# Closed-Loop Estimation of Neurostimulation Strength-Duration Curve Using Fisher Information Optimization and Comparison With Uniform and Random Methods

**DOI:** 10.1101/2023.10.19.563097

**Authors:** Seyed Mohammad Mahdi Alavi

## Abstract

**Background:** Strength-duration (SD) curve, rheobase and chronaxie parameters provide insights about the interdependence between stimulus strength and stimulus duration (or pulse width), and the neural activation dynamics such as the membrane time constant, which are useful for diagnostics and therapeutic applications. The existing SD curve estimation methods are based on open-loop uniform and/or random selection of the pulse widths.

**Objective:** To develop a method for closed-loop estimation of the SD curve.

**Method:** In the proposed method, after the selection of each pulse width through Fisher information matrix (FIM) optimization, the corresponding motor threshold (MT) is computed, the SD curve estimation is updated, and the process continues until satisfaction of a stopping rule based on the successive convergence of the SD curve parameters. The results are compared with various uniform methods where pulse widths are chosen in ascending, descending and random orders, and with methods with two and all non-uniform random pulse widths.

**Results:** 160 simulation cases were run. The FIM method satisfied the stopping rule in 144 runs, and estimated the rheobase (chronaxie in parenthesis) with an average absolute relative error (ARE) of 1.73% (2.46%), with an average of 82 samples. At this point, methods with two and all random pulse widths, and uniform methods with descending, ascending and random orders led to 5.66% (20.27%), 2.15% (4.51%), 8.57% (54.96%), 3.52% (5.45%), and 2.19% (4.40%) AREs, which are greater than that achieved through the FIM method. In all 160 runs, The FIM method has chosen the minimum and maximum pulse widths as the optimal pulse widths.

**Conclusions:** The SD curve is identifiable by acquiring the SD data from the minimum and maximum pulse widths achieved through the FIM optimization. The SD data at random or uniform pulse widths from only the vertical area or lower plateau of the curve might not result in satisfactory estimation.

**Significance:** This paper provides insights about pulse widths selection in closed-loop and open-loop SD curve estimation methods.

## I. Introduction

Activation of a neural tissue depends not only on stimulus strength, but also on stimulus duration. By plotting motor threshold (MT) versus stimulus duration, strength-duration (SD) curve provides insights about the interdependence between stimulus strength and duration in activating a nerve, [1], [2], [3]. The SD curve is typically represented by a rectangular hyperbola, shown in Fig. 1.

**Fig. 1.**
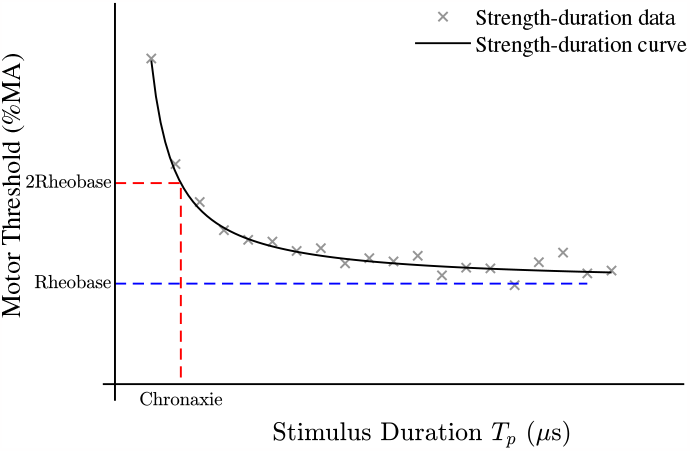
Strength-Duration Curve.

MT is defined as a minimum stimulus strength which results in minimal neural activation. It is detected visually or through electromyography (EMG). In the visual technique, MT is defined as the minimal pulse intensity that leads to a visible muscle twitch or contraction by stimulating the motor cortex area (M1) of that muscle, [4]. In the EMG-based techniques, MT is defined as the minimal pulse intensity that leads to a predefined motor evoked potential (MEP) on the target muscle [5], [6]. Conventionally, MT is measured by using relative frequency methods by administering sequential pulses with intensities in ascending or descending orders, until at least 50 *µ*V MEP is observed in 50% of 10 to 20 consecutive stimuli [5] and [6]. In more advanced techniques, MT is computed using maximum-likelihood (ML), maximum a-posteriori (MAP), or Bayesian inference theories, [7]–[10]. The visual MT detection technique is more convenient to perform, and it requires no expertise in EMG. However, studies confirm that the EMG-based methods detect MT more accurate and faster with fewer stimuli, compared to the visual techniques, [7], [11]–[15].

Each point on the SD curve determines the MT value and stimulus duration to elicit a minimal muscle contraction or action potential in the target nerve or muscle.

Rheobase on the SD curve represents the MT value for infinite stimulus duration. When the stimulus strength is below the rheobase, stimulation is ineffective even if the stimulus duration is very long.

Chronaxie is another point on the SD curve, which represents a pulse duration at which the MT value becomes twice the rheobase. Chronaxie is inversely proportional to excitability. As chronaxies varies among neural structures, stimulus pulse duration is an important element for selective stimulation and electrophysiological classification. It was experimentally shown that the intracellular to extracellular chronaxie ratio is two or higher, depending on the cell properties, distance and type of electrodes [16]. Studies also confirm the relationship between chronaxie and membrane time constant. In [17], the chronaxie and membrane time constant is assumed to be almost equal. In [16], [18], it is suggested that the chronaxie is often about 0.7 times the membrane time constant for near-rectangular pulses.

Thus, SD curve is a powerful tool for studying the nerve system, diagnostics and therapeutic applications. The results in [19] imply that the strength-duration curve, the chronaxie and the rheobase parameters could be useful in assessing spinal cord function. In [20], a SD-curve-based technique was developed to find the most adequate pulse widths in rechargeable neurostimulation systems to obtain the largest coverage of the painful area, the most comfortable paresthesia, and the greatest patient satisfaction. In [21], SD curve was used to calculate and compare time constants of motor cortex structures activated by current pulses oriented posterior–anterior or anterior–posterior across the central sulcus. In [22], the excitability of the common fibular nerve in subjects with and without ankle injuries was studied using the SD curve. In [23], SD curve was used to analyze the relationship between electrophysiologic excitability and morphology of the radial nerve in patients with unilateral chronic lateral epicondylalgia. By using SD curve, it was shown that, for the treatment of chronic pain during spinal cord stimulation, smaller dorsal column fibers can only be activated with sufficiently large stimulus pulses [24]. In [25], it was shown that the cortical SD curve properties also depend on the pulse shape and pulse width selection in transcranial magnetic stimulation (TMS). The SD results through single pulse electrical stimulation indicated that high current and short pulse width stimulations elicit strong and consistent responses while minimizing charge, [26].

For the SD curve estimation, MT values for a set of pulse widths are firstly computed. The SD curve is then estimated by fitting to the data set, as shown in Fig. 1. Weiss [27] and Lapicque [28] SD curve models have widely been utilized in the literature, [1], [17], [29]–[31]. Weiss proposed the hyperbolic form of the SD curve, *I* = *b*(1 + *c/d*), where *I* is the threshold current, *b* is the rheobase current, *c* is the chronaxie, and *d* is the pulse width, and Lapicque proposed the exponential SD curve in the form of *I* = *b/*(1 −*exp*(− *d/c*)), [32].

### A. The contributions of this paper

In the existing literature, the pulse widths are chosen randomly and/or uniformly distributed, in an open-loop framework, prior to the experiments [1], [17], [26], [31], [33]. There is no systematic method or result to determine how many data points are enough to estimate a SD curve with a desired level of accuracy.

This paper proposes a method for closed-loop estimation of the SD curve using Fisher information matrix (FIM) optimization. In the proposed method, the pulse widths are chosen sequentially, by using the prior data and FIM optimization. After the selection of each pulse width, the corresponding MT is computed, the SD curve estimation is updated, and the process continues until satisfaction of a stopping rule based on the successive convergence of the SD curve parameters.

The effectiveness of the proposed FIM-based SD curve estimation method is evaluated and discussed through extensive comparison with the uniform and random based estimation methods on 160 simulation cases.

In [31], [34]–[36], the FIM optimization was developed for closed-loop estimation of the neural input-output curve and activation dynamics including the membrane time constant and coupling gain. No paper or study has been published on the FIM-based closed-loop SD curve estimation, and comparison with the uniform and random methods.

### B. The structure of the paper

This paper is organized as follows. The SD curve model and estimation problem are described in Section II. Section III elaborates the FIM-based SD curve estimation method. The simulation results are discussed in Section IV.

## II. Strength-Duration Curve Model and Estimation Problem

Without loss of generality, this paper focuses on the Weiss SD curve model, given by:

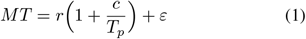

where, *MT* is motor threshold, *r* is rheobase, *c* is chronaxie, *T*_*p*_ is stimulus duration, and *ε* represents the MT variability. Development to the Lapicque model is similar and straight-forward.

The problem is to estimate the parameter vector

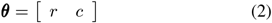

in a closed-loop system, until a satisfactory level of estimation is achieved.

Vectors and matrices are denoted by bold in this paper.

## III. Estimation Using Fisher Information Matrix Optimization

The computational steps of the FIM-based SD curve estimation are given in Algorithm 1.

In the initialization step, two random pulse widths, 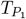 and 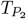, are randomly chosen, and their corresponding MTs, *MT*_1_ and *MT*_2_, are computed by using one of the methods discussed in Section I.

An initial estimation of the parameter vector is obtained by fitting the SD curve model (1) to the initial data set 𝒟 = 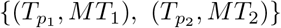.

### Algorithm 1: Strength-duration (SD) curve estimation by using Fisher information optimization

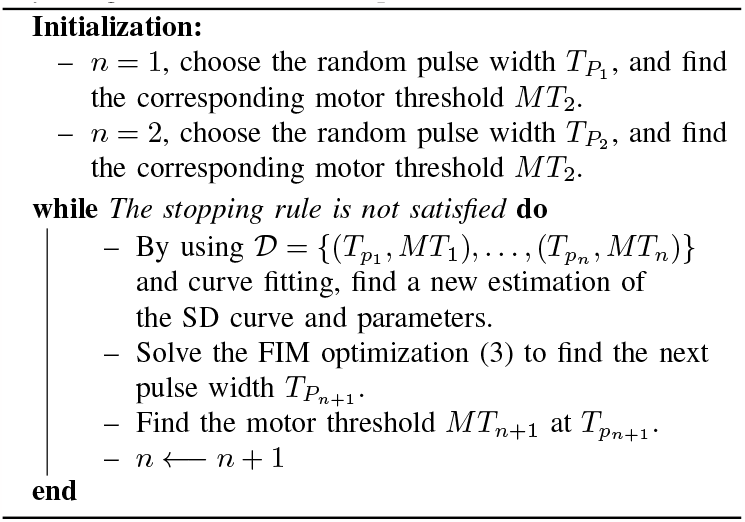

By using the prior data, the next pulse width, 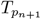, is calculated by optimizing the (*n* + 1)–th FIM in a closed-loop system as follows:

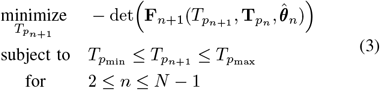

where, *T*_*pn*_ = [*T*_*p*1_ … *T*_*pn*_] is the prior pulse widths vector, 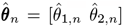 is the estimated parameter vector after the *n* −th pulse width and MT calculation, 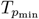 and 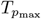 are the predefined minimum and maximum pulse widths, and *N* is the predefined maximum number of data points.

The (*n* + 1)*−*th FIM is computed recursively as follows:

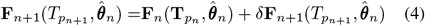

where,

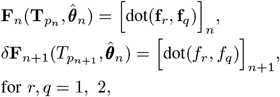

and dot(**x, y**) returns the scalar product of any vectors **x** and **y**.

For the SD curve (1), **f**_*i*_ and *f*_*i*_, *i* = 1 2 are given by:

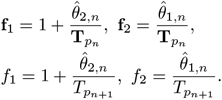

After calculation of the next pulse width 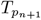, the corresponding MT is calculated, and the parameter estimation is updated by fitting the SD curve model to the data set of

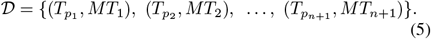

Finding the next pulse width and MT, and updating the data set and parameter estimation are repeated until a stopping rule is satisfied.

The stopping rule is defined as the converge of the estimation parameters

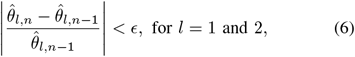

for a predefined successive times *T* = 1, 2, 3, .… The parameters 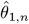 and 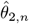 denote the estimations of the rehobase and chronaxie at the *n*-th sample, respectively, and *ϵ* is the convergence tolerance. The smaller the *ϵ* value and the larger the *T* value, the more the accurate estimation; however, more SD data will be needed.

A ‘force stop’ is defined to terminate the estimation process if the stopping rule is not satisfied at the maximum sample number *N* .

‘Termination sample’, *n*_*s*_, is defined as the sample number at which the estimation process stops, thus, *n*_*s*_ ≤ *N* .

‘Successful termination’, *n*_*ss*_, means that the stopping rule is satisfied prior to the force stop, i.e., *n*_*ss*_ = *n*_*s*_ < *N* .

The termination sample and successful termination sample of the FIM-based SD curve are denoted by 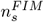 and 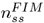, respectively.

## IV. Results and Discussions

The effectiveness of the proposed FIM-based SD curve estimation method is evaluated through 160 simulation runs in Matlab R2023a (The MathWorks, Inc.).

For each simulation run, a true model is generated using (1), with true parameters randomly chosen within the following range:

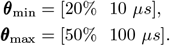

The objective is to estimate the true SD curve and parameters, and compare the results.

The MT variability *ε* is assumed to be a zero-mean white noise with a random variance arbitrarily chosen between 2% and 6%.

The stopping rule is arbitrarily defined to satisfy the convergence criterion (6) with *ϵ* = 0.001 for *T* = 5 successive times. As discussed in [31], [34]–[36], there is a trade-off between the estimation accuracy (i.e., *ϵ* and *T* values) and the number of samples in a successful termination. Reducing the convergence tolerance and increasing the successive times parameter would improve the estimation, however, more SD data are needed to meet the stopping rule. The force stop is set to happen at *N* = 200.

The trust-region curve fitting algorithm with lower and upper limits ***θ***_min_ and ***θ***_max_ is used for updating the estimation in all simulation runs.

The FIM optimization (3) is solved by using the ‘fmincon’ and global search interior-point algorithms with a random initial guess. The lower and upper limits of the pulse width are set to

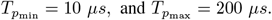

For comparative studies, the SD curve is also estimated using uniform and random sampling methods. In the uniform-based methods, SD curve is estimated with *N* pulse widths uniformly distributed between 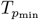 and 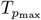. Three different uniform-based strategies are simulated, where pulse widths are chosen in ascending, descending and random orders.

Apart from the random-order uniform method, two other random-based methods are simulated with two or all random pulse widths. In the two random pulse widths method, the SD curve is estimated by using SD data at only two pulse widths, which are chosen randomly between 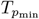 and 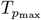 . The SD data are computed at these two pulse widths recursively until termination. In the all random pulse widths method, the SD curve is estimated by different pulse widths, all of them chosen randomly between 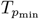 and 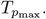

In all methods, the sequential estimation approach is implemented, which means that, the SD curve estimation is updated and the stopping rule is checked after each stimulus. For consistency, the same curve fitting method and stopping rule as in the FIM-based method are utilized.

Termination samples of all runs and methods are recorded and compared.

For quantitative analysis, the absolute relative error (ARE) of the estimated parameter vector, **e**_***θ***,*n*_, is computed at every sample as follows:

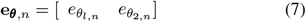

where, *n* is the sample number, and

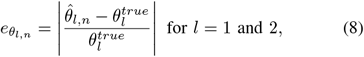

where, 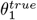 and 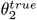 are true rheobase and chronaxie values, and 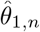 and 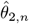 are their estimations at the *n*-th sample.

The ARE of each estimation method is shown by superscript, e.g., 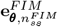 and 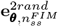 denote the estimation AREs for the FIM and two random pulse widths methods at the successful termination of the FIM method, respectively.

The average AREs over 160 runs is also computed and discussed for all methods.

### A. A representative run

Fig. 2 shows the SD curve of a representative run, with the following true parameters:

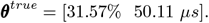

**Fig. 2.**
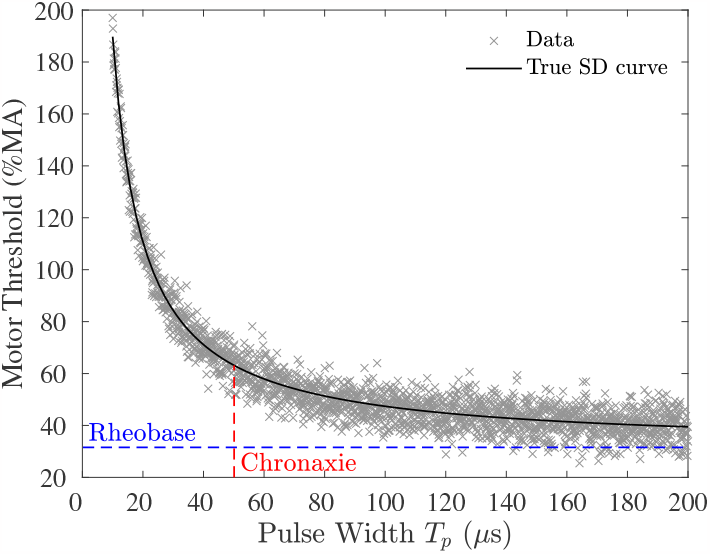
Sample simulation run; the SD curve and parameters, and variability of the motor threshold (MT) measurement with respect to the pulse width.

The variability of the MT values is generated by a random zero-mean noise with the variance of 5.30%.

For this case study, the FIM SD curve estimation is successfully terminated at 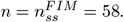

Fig. 3 shows the samples and estimated SD curves for all methods at 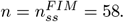

**Fig. 3.**
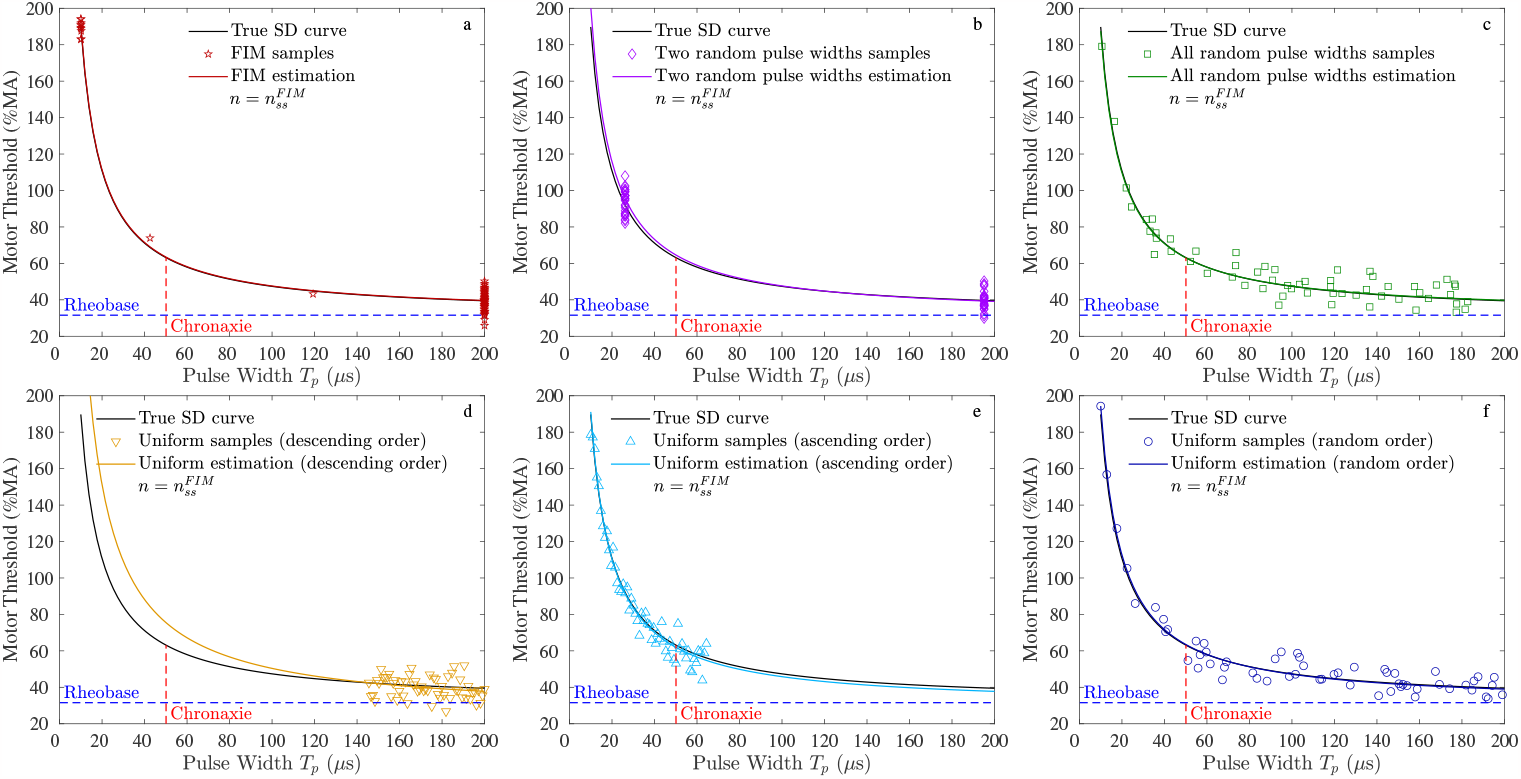
Sample simulation run; samples and estimated SD curves at 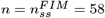, where the FIM method successfully terminated. a) The FIM method. b) The two pulse widths method. c) The all random pulse widths method. The uniform method, administered in d) descending order, e) ascending order, and f) random order.

Fig. 4 shows the samples and estimated SD curves at *n* = *N* = 200.

**Fig. 4.**
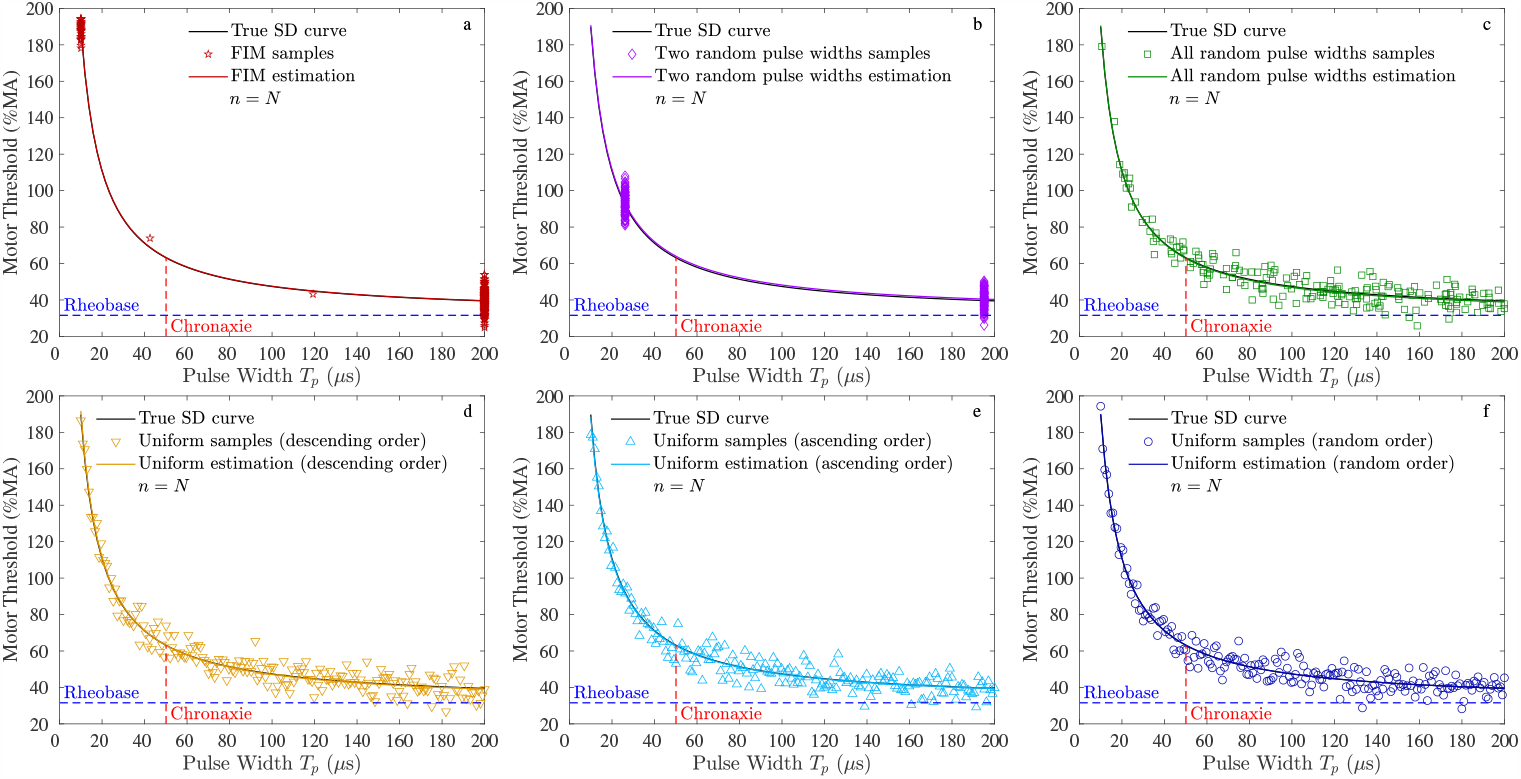
Sample simulation run; samples and estimated SD curves at *n* = *N* = 200. a) The FIM method. b) The two pulse widths method. c) The all random pulse widths method. The uniform method, administered in d) descending order, e) ascending order, and f) random order.

Fig. 5 shows the rheobase and chronaxie estimated at every sample.

**Fig. 5.**
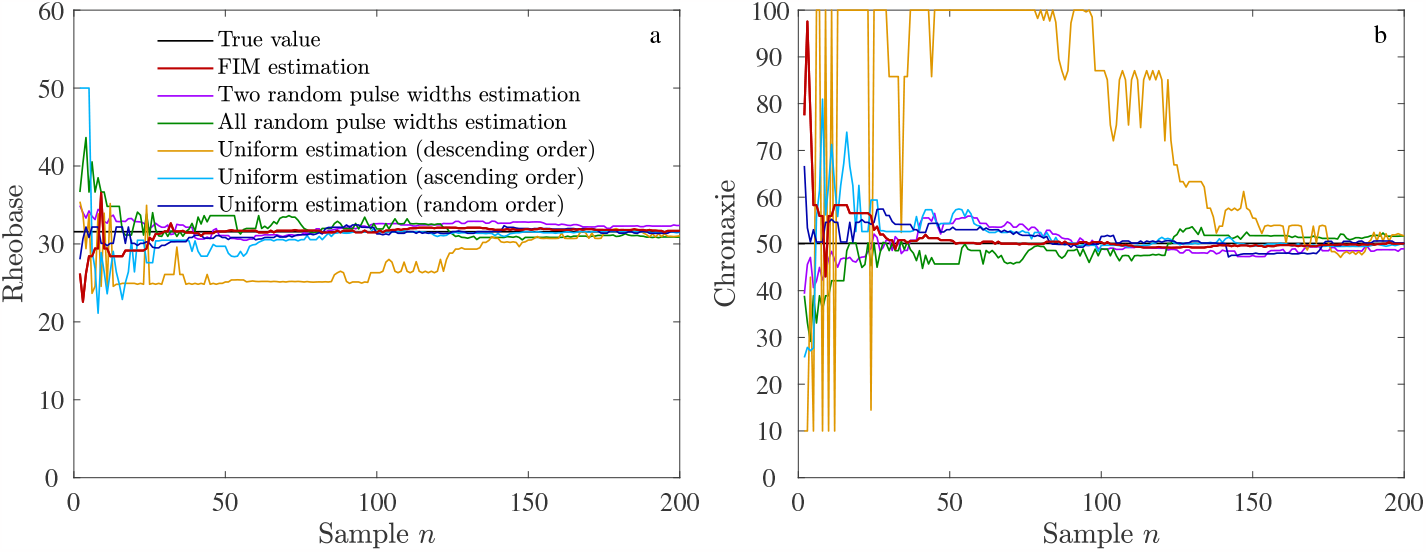
Sample simulation run; estimated values of the rheobase and chronaxie parameters.

As seen in Fig. 3-a and Fig. 4-a, the FIM method selects 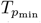 and 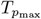 as the optimal pulse widths, and estimates the SD curve with AREs of 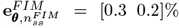 and 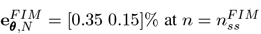 and *n* = *N*, respectively. The two middle data points are the random initial pulse widths, which could be chosen from 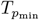 and/or 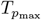. Thus, it can be induced that data at two pulse widths would be enough for the SD curve estimation. This is aligned with the structural identifiability analysis [37], which says a nonlinear equation with two unknown parameters is identifiable by using two data points. However, acquiring multiple data and sequential updating of the estimation help reduce the estimation error in the presence of MT variability or measurement noise. This is so-called the practical identifiability [38], and as illustrated in this example, the FIM SD curve estimation method efficiently performs it.

In order to illustrate that the FIM method selects the optimal pulse widths, the SD curve is also estimated with the data acquired only at two pulse widths chosen randomly between 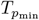 and 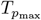, as shown in Fig. 3-b and Fig. 4-b. It is seen that the true and estimated SD curves are not matched at 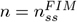, and more data points are required to achieve a better fit, as shown in 4-b. The AREs at 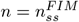 and *n* = *N* are 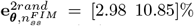 and 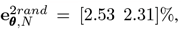 respectively, which are significantly larger than those obtained through the FIM method.

In Fig. 4-b, the two random pulse widths are chosen from the two sides of the chronaxie. Fig. 6 shows the results when the two random pulse widths are chosen from one side of the chronaxie, from the vertical area close to 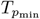 or from the lower plateau close to 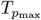. It is observed that the estimation AREs increase, compared to the FIM method and when the two pulse widths are chosen from the both sides of the chronaxie. If the pulse widths are chosen close to 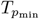, the SD curve around 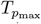 might not be estimated well, and vice versa. The estimation error is significant when the two pulse widths are chosen from the lower plateau, even with *n* = *N* = 200 samples as shown in Fig. 6-d. The SD curve estimations in Figures 6-a,c result in the AREs of 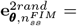 [7.87 11.23]% and 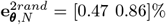 at 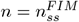 and *n* = *N*, respectively. The SD curve estimations in Figures 6-b,d result in the AREs of 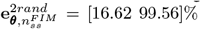 and 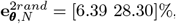, respectively.

**Fig. 6.**
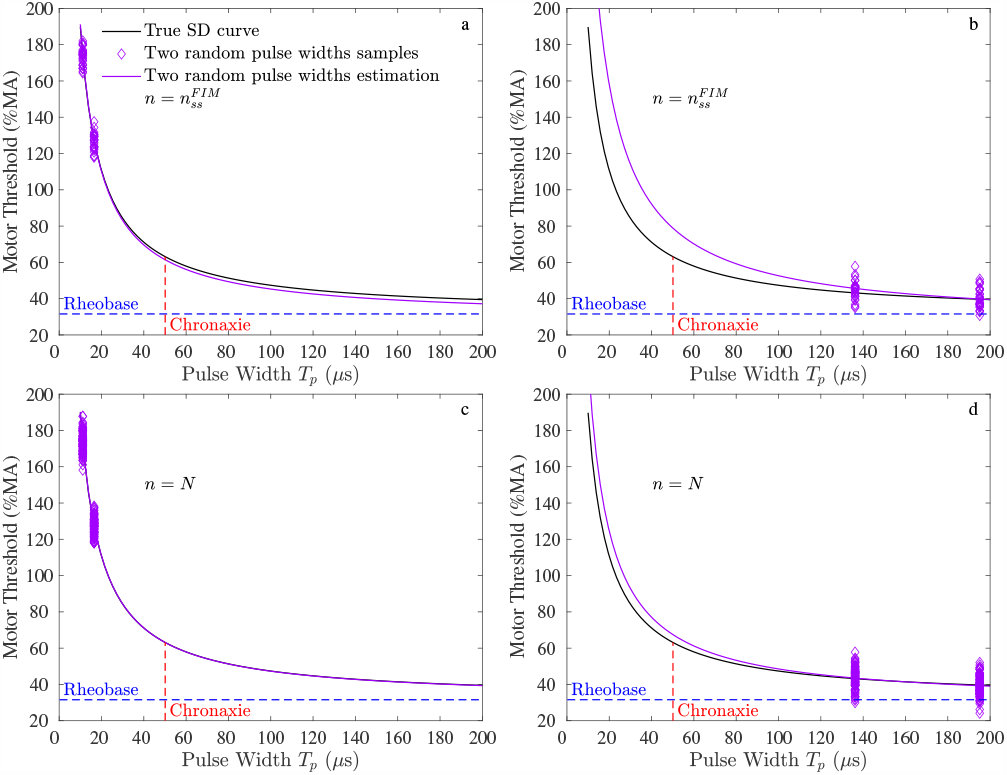
Sample simulation run; samples and estimated SD curves with two random pulse widths chosen close to 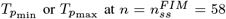 (a and b) and at *n* = *N* (c and d).

Only the two random pulse widths from both sides of the chronaxie results in a successful termination at 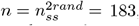, and force stop happens when the two random pulse widths are chosen from one side of the chronaxie.

Fig. 3-c and Fig. 4-c show the SD curve estimations with all random pulse widths at 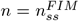 and *n* = *N*, respectively. The estimation results are satisfactory, with the estimation AREs of 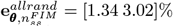 and 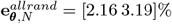 at 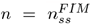 and *n* = *N*, respectively. The successful termination happens at 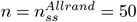 with the all random pulse widths method.

Figures 3-d,e,f show the SD curve estimations with uniformly distributed pulse widths in descending, ascending, and random orders at 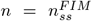, respectively. Their data at *n* = *N* are shown in Figures 4-d,e,f. The estimation AREs at 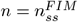 and *n* = *N* are as follows:

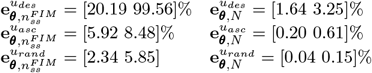

At 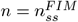, only the random-order uniform method results is a satisfactory estimation as shown in Fig. 3-f. The estimation errors of the uniform methods with ascending and specially descending orders are considerable as shown in Fig. 3-d and Fig. 3-e. The estimated rheobase and chronaxie shown in Fig. 5 confirm this result. This is consistent with the afore-mentioned finding that SD data only from the vertical area or lower plateau might not lead to a satisfactory estimation.

At *n* = *N* = 200, all uniform methods result in satisfactory estimations. However, the major issue of the uniform methods should be re-called that, the optimal value of *N* is not known prior to the experiment, and *N* is chosen randomly based on the trials and errors in the literature.

The successful termination happens at 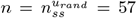 and 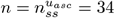 for the uniform method based on random and ascending orders. The uniform method with descending order is terminated with force stop at *n* = *N* = 200.

### B. Multiple runs

The following results are achieved through 160 simulation runs.

144 (90%) of the FIM runs, 139 (86.88%) of the runs with all random pulse widths, 53 (33.12%) of the runs with two random pulse widths, and 76 (47.5%), 129 (80.63%), and 138 (85.6%) of the uniform runs with descending, ascending, and random orders result in successful termination with an average of 82, 109, 79, 100, 87 and 87 samples, respectively.

Fig. 7 shows the average AREs of the rheobase and chronaxie estimations at the successful termination samples for all methods.

**Fig. 7.**
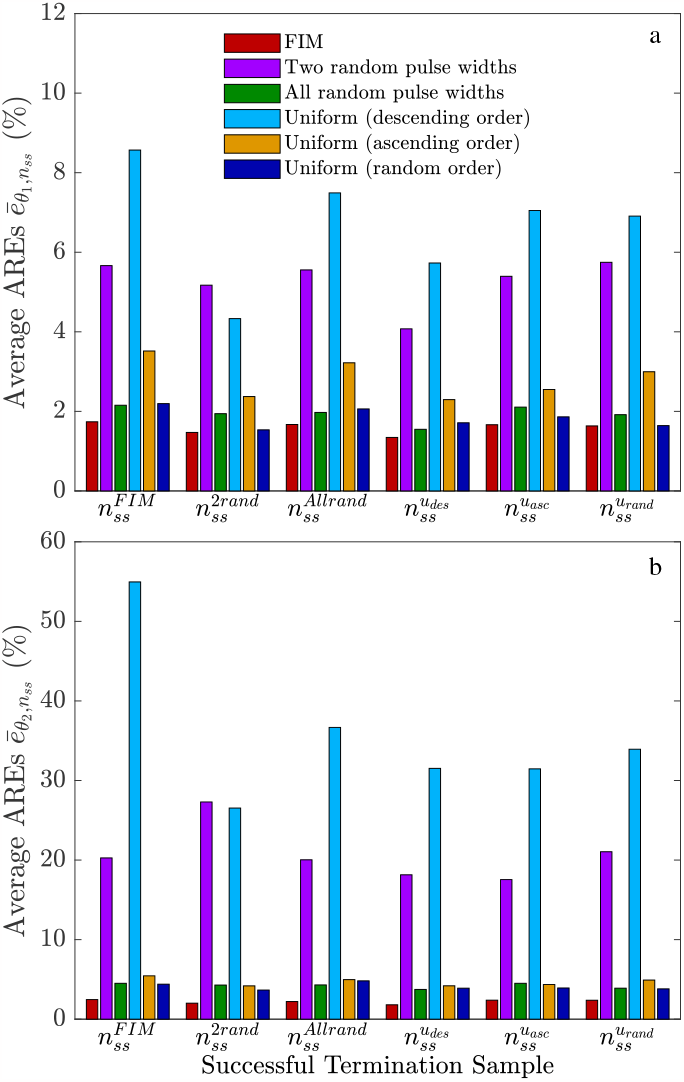
Average relative errors (AREs) of the rheobase and chronaxie estimations in 160 runs.

At the FIM method’s successful termination, i.e. 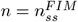, the rheobase (chronaxie in parenthesis) is estimated with the average AREs of 1.73% (2.46%), 5.66% (20.27%), 2.15% (4.51), 8.57% (54.96%), 3.52% (5.45%), and 2.19% (4.40%) with the FIM method, methods with two and all random pulse widths, and uniform methods with descending, ascending and random orders, respectively.

At their own successful terminations, i.e., 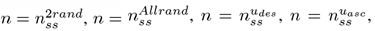, and 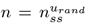, methods with two and all random pulse widths, and uniform methods with descending, ascending and random orders estimated the the rheobase (chronaxie in parenthesis) with the average AREs of 1.73% (2.46%), 5.17% (27.30%), 1.97% (4.30%), 5.73% (31.52%), 2.55% (4.36%), and 1.64% (3.81%), respectively.

Overall, the following key points are deduced from the AREs’ results: 1) The rheobase parameter is estimated with a better resolution in average, compared to the chronaxie parameter. 2) The FIM estimation method results in the smallest average AREs not only at 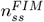, but also in the successful terminations of other methods. In the next ranks, the random order uniform, and all random pulse widths methods perform better. 3) As discussed earlier, the uniform method with descending order as well as the method with two random pulse widths lead to the worst performance in terms of the estimation AREs.

In all of the 160 simulation runs, the FIM method selects 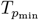 and 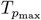 as the optimal pulse widths, and the true SD curve is accurately estimated as illustrated above. By selecting the initial pulse widths from 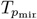 and/or 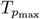, there is no need to solve the optimization problem (3), which will reduce the computational burden.

## V. Conclusions

The results of this study confirm that the FIM SD curve estimation method outperforms the uniform and random methods, by acquiring the SD data at only two pulse widths. It was illustrated that the location of two pulse widths is important. The results confirm that sampling from the minimum and maximum pulse widths in the FIM method is the optimal strategy compared to other estimation methods based on the two random pulse widths from one or both sides of the chronaxie. It was further shown that acquiring samples only from the vertical area or lower plateau might not result in a satisfactory estimation. After the FIM method, the results of the randomly (uniformly or non-uniformly distributed) pulse widths resulted in a better SD curve estimation performance.

